# Nod2 protects remote small intestinal sites in case of colonic inflammation

**DOI:** 10.1101/510933

**Authors:** Ziad Al Nabhani, Dominique Berrebi, Christine Martinez-Vinson, Nicolas Montcuquet, Gilles Dietrich, Gurminder Singh, Jerrold R. Turner, Chrystele Madre, Maryline Roy, Eric Ogier-Denis, Monique Dussaillant, Nadine Cerf-Bensussan, Habib Zouali, Camille Jung, Fanny Daniel, Frédérick Barreau, Jean-Pierre Hugot

**Affiliations:** Laboratoire d’excellence Inflamex, Université Paris-Diderot Sorbonne Paris-Cité, UMR 1149, F-75018 Paris, France; INSERM, UMR 1149, F-75018 Paris, France; Assistance Publique Hôpitaux de Paris, services des maladies digestives et respiratoires de l’enfant et service d’anatomie pathologique, Hôpital Robert Debré, F-75019 Paris, France; INSERM, UMR 989, F-75015 Paris, France; Université Paris Descartes - Sorbonne Paris Cité, Institut IMAGINE, F-75015 Paris, France; IRSD, Université de Toulouse, INSERM, INRA, ENVT, UPS, Toulouse, France; Department of Pathology, Brigham and Women’s Hospital, Harvard Medical School, Boston, Massachusetts; Fondation Jean-Dausset CEPH, F-75011 Paris, France

**Keywords:** Nod2, intestinal permeability, Crohn’s disease, gut barrier, myosin light chain kinase, CD4^+^ T cells

## Abstract

*NOD2* mutations are key risk factors for Crohn’s disease (CD). NOD2 contributes to intestinal homeostasis by regulating innate and adaptive immunity together with intestinal epithelial function. However, the roles of NOD2 during gut inflammation is not known. We initially observed that *NOD2* expression was increased in epithelial cells remote from inflamed areas in CD patients. To explore this finding, *Nod2* mRNA expression, inflammation and gut permeability were examined in the small bowel of wild-type (WT), *Nod2* knockout and *Nod2* mutant mice after rectal instillation of 2,4,6-trinitrobenzene sulfonic acid (TNBS). In WT mice, Nod2 upregulation remote to rectal injury was associated with pro-inflammatory cytokine expression, recirculating CD4^+^ T-cells, increased paracellular permeability and myosin like chain kinase activity. *Nod2* knockout or mutation led to duodenitis and ileitis demonstrating the remote protective role of Nod2. Bone morrow stem cell (BMSC) transplantations indicated that the small intestinal inflammation was due to NOD2 loss in both hematopoietic and non-hematopoietic compartments. As a whole, WT but not mutant NOD2 prevents disease extension at sites remote from the initial intestinal injury.

## Introduction

Crohn’s Disease (CD) is an inflammatory bowel disease (IBD) that can affect any part of the entire gastrointestinal tract. Genetic and epidemiological studies indicate that CD is a complex, multifactorial disorder. Interplay between genetics and the environment promotes development of gut abnormalities of autophagy, reticulum endoplasmic stress, innate and adaptive immune responses, Th-1 and Th-17 polarization, intestinal barrier dysfunction and microbial dysbiosis.^1–3^

Nucleotide oligomerization domain 2 (*NOD2*, also known as NLR-C2 and CARD15) is the most prominent susceptibility gene for CD.^4, 5^ One-third to one-half of CD patients have one or more *NOD2* mutations.^6^ Wild-type NOD2 is activated by muramyl dipeptides (MDP) which are components of the bacterial cell wall,^7^ but CD-associated *NOD2* mutations prevent MDP responses.^8^ CD can therefore be considered as an immune deficiency with insufficient responses to bacteria. Nevertheless, the exact mechanism by which *NOD2* mutations contribute to CD pathogenesis remains a matter of debate.^9–12^

*NOD2* regulates innate and adaptive immunity and intestinal permeability to maintain intestinal homeostasis.^13–16^ Indeed, *Nod2* ablation in mice leads to an increased bacterial translocation across the small intestinal epithelium and excessive inflammatory cytokine secretion.^14, 15^ This reflects impaired crosstalk between inflammatory cytokine-secreting CD4^+^ T-cells and epithelial cells that express myosin light chain kinase (MLCK).^15, 17^ Similarly, increased MLCK activity^18^ and CD4^+^ T-numbers have been observed in the intestinal mucosa of CD patients,^19, 20^ and mouse models show that genetic activation of epithelial MLCK induces increases in mucosal CD4^+^ T-numbers.^21^ Anti-TNF-α antibody treatment restores the intestinal barrier in CD patients.^22^ Impaired epithelial barrier function may therefore be an early event in CD lesions progression.

Here, we show that NOD2 expression in CD patients is not only increased in inflammatory lesions but also at sites remote from injury. To define the mechanisms and impact of this upregulation, we explored the remote consequences on the small intestinal mucosa of a limited rectal injury induced by 2,4,6-trinitrobenzene sulfonic acid (TNBS).

## Results

### Epithelial NOD2 expression is increased in uninflamed mucosa of CD patients

In CD patients, epithelial *NOD2* expression is increased in mildly inflamed areas of the digestive tract, remote from sites of injury.^23^ To confirm NOD2 upregulation in remote areas, we examined expression in ileal and/or cecal biopsies from 17 treatment-naïve pediatric CD cases and five non-inflammatory controls. Although nine CD patients had heterozygous mutations in *NOD2* (1007fs n=3, R702W n=5 and R373C n=1), no histological differences were seen between patients with wild type or mutant *NOD2*. Immunostains using two different antibodies showed that NOD2 was weakly expressed by surface enterocytes and rare mononuclear cells immediately below the epithelium in control ileum (Figure 1*A*). In contrast, NOD2 expression was increased in ileum from CD patients (Figure 1 *B* and *C*). While NOD2 expression was upregulated in lamina propria mononuclear cells within inflammatory areas, the most prominent increases were in surface and glandular epithelial cells outside of inflammatory lesions (Figure 1*B* and *C*). Analysis of cecal biopsies gave similar results (Figure 1*D-F*).

**Figure 1:**
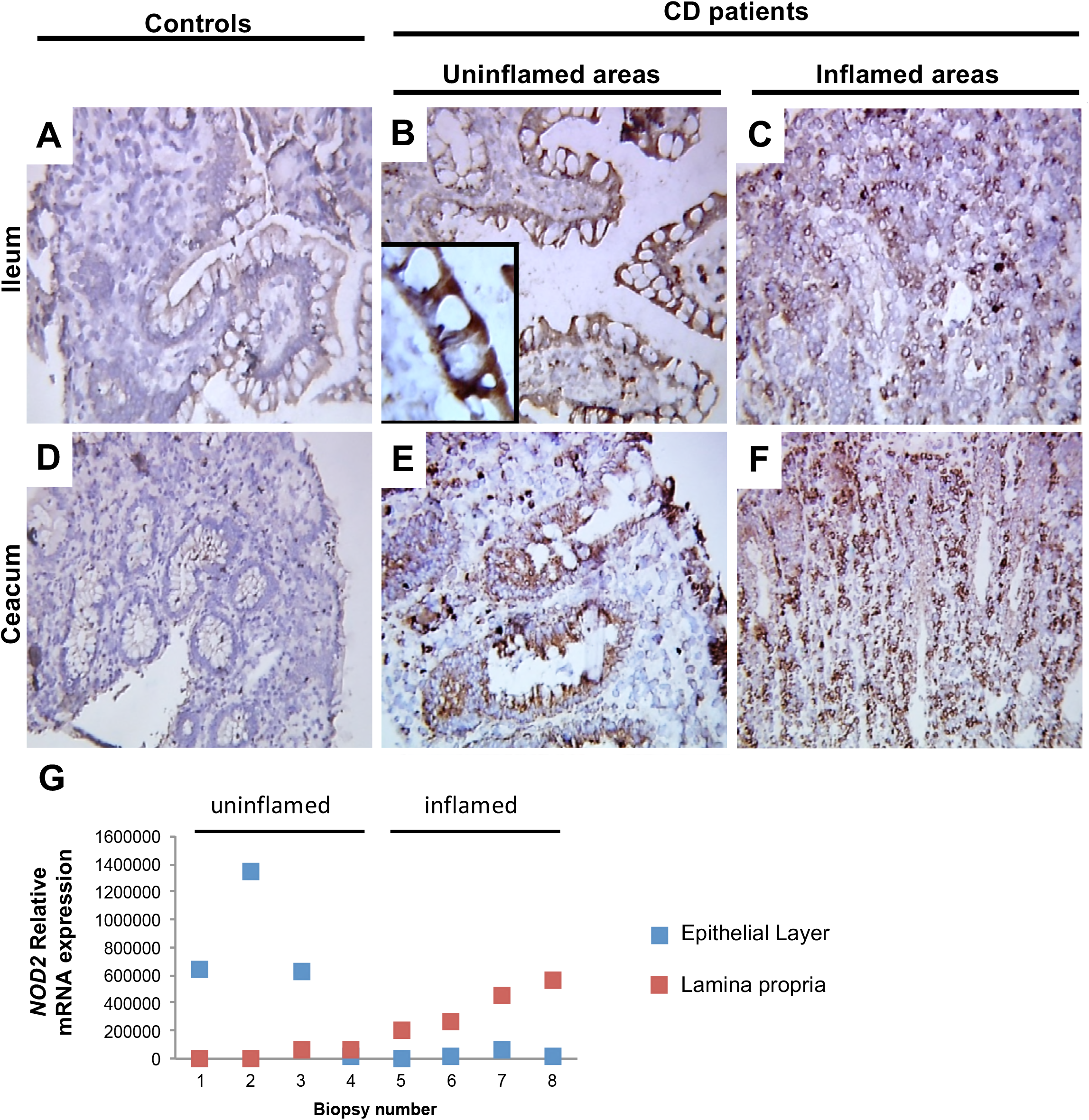
NOD2 expression in increased in intestinal mucosa of Crohn disease patients. *(A-F)* Biopsies from controls *(A* and *D)* and naïve pediatric CD patients *(B, C, E* and *F)* were immunostained with anti-NOD2 antibodies. Ileal (A-C) or cecal (D-F) biopsies were obtained from inflamed *(C* and *F)* or uninflamed areas *(B* and *E)*. Data shown are representative of 5 controls and 17 CD patients. *(G)* Number of *NOD2* mRNA copies were normalized by the expression of Abelson gene and expressed as arbitrary units. mRNA levels were calculated for the epithelial monolayer (in blue) and the lamina propria (in red) from the same biopsy after laser microdissection. Biopsies were obtained from inflamed or uninflamed intestinal areas and referenced by an arbitrary number.

We then determined *NOD2* mRNA expression in the epithelial and lamina propria compartments by qPCR after laser microdissection of biopsies from 8 patients. We observed that *NOD2* mRNA expression was inversely correlated in the epithelial and lamina propria compartments of the same biopsy (Figure 1*G*). In the lamina propria, the average *NOD2* copy number was 43.1 in controls (normalized arbitrary units). In CD patients, similar values (43.6) were observed in uninflamed areas whereas *NOD2* expression was increased a 5-fold (205.907) in the inflamed ileum. On the contrary, the mean values were 4.91 in epithelial cells of controls and 4.6 in inflamed ileal areas but a 100-fold increase in *NOD2* expression (660) was detected in uninflamed ileum. Noteworthy, normal Paneth cells had low *NOD2* expression in controls (1.43). This expression was increased by inflammation and in heterotopic colonic Paneth cells but *NOD2* was mostly expressed by enterocytes. We thus concluded that *NOD2* expression is markedly increased in epithelial cells distant from inflammatory lesions in CD patients.

### Gut injury leads to epithelial Nod2 expression and cytokine production at remote sites via CD4^+^ T-cell activation

To confirm the expression of epithelial Nod2 in healthy areas distant from intestinal lesions in a mouse model, we treated *Nod2* wild-type (*Nod2*^WT^) mice by an intra-rectal administration of TNBS. Instillation of TNBS in mice is known to induce a severe inflammation in the distal colon.^14^ Interestingly, TNBS has also been shown to alter the biochemical activity of brush border enzymes (sucrase isomaltase and aminopeptidase), mucins and cytokines levels in the small bowel (*i.e*. at a significant distance from the gut injury) without histological lesions.^24^ Three days after instillation, mice were sacrificed and the severity of inflammation was assessed (Figure 2). In the distal colon, TNBS administration induced a robust inflammation as evidenced by decreased body weight, increased disease activity index (DAI), reduced colon length and high macroscopic Wallace damage scores (Figure 2*A-D*). Consistent with this phenotype, expression levels of TNF-α, IFN-γ and IL-12 were increased (Figure 2*E*) at the site of colonic inflammation.

**Figure 2:**
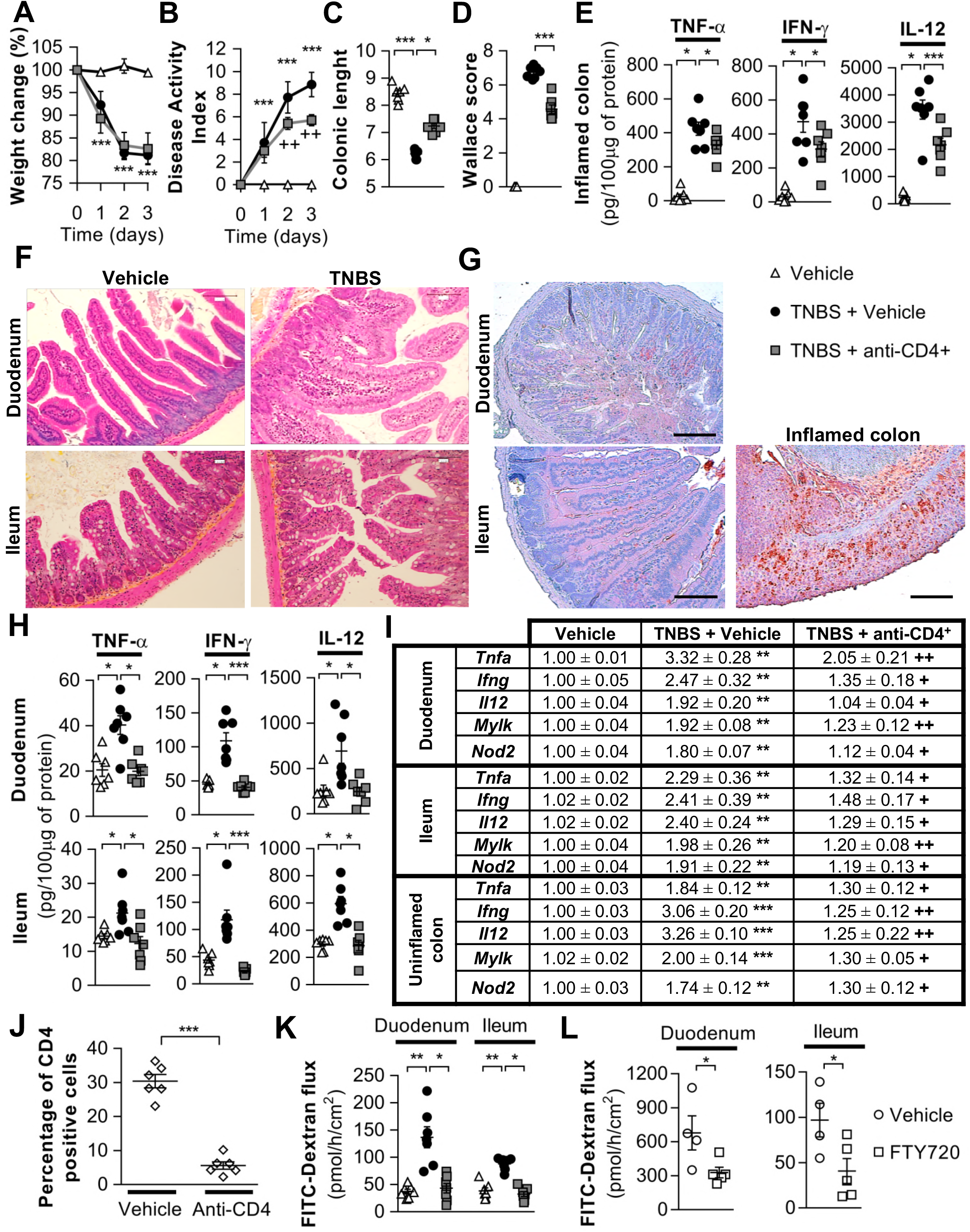
Remote gut barrier dysfunctions of the small intestine are mediated by recirculating CD4^+^ T-cells. *(A-L)* C57BL/6 wild-type mice (*Nod2*^WT^) were instilled intra-rectally with TNBS. Vehicle control group was challenged by PBS-Ethanol. Mice were treated with anti-CD4^+^ antibodies or FTY720, an inhibitor of T-cell recirculation where indicated. 3 days after the instillation, the intensity of the colitis was monitored with the following parameters: *(A)* Weight loss; *(B)* Disease activity index; *(C)* Colonic length (cm); *(D)* Colonic macroscopic score (Wallace score); *(E)* Pro-inflammatory cytokine levels in inflamed colon. In parallel, the duodenum and ileum were explored by microscopic examination after (F) hematoxylin-eosin staining and (G) Myeloperoxidase (MPO) immunostaining (colonic MPO expression in a mouse treated with DSS 3/ for 7 days is shown as a positive inflammatory control); H) Protein levels of pro-inflammatory cytokines; (I) mRNA expressions of pro-inflammatory cytokines, *Mlyk* and *Nod2;* (J) levels of CD4^+^ T-cells in ileal Peyer’s patches after CD4^+^ depletion with anti-CD4^+^ antibodies; (K-L) Paracellular permeability assessed by Ussing chamber experiments. Original magnification, X20. Scale bars: 100μM. (Each point = one mice; mean±s.e.m; 3 independent experiments; ^*^P<0.05, ^*^P<0.01 and ^***^P<0.001 vs. vehicle control group or indicated group; ^++^P<0.01 vs. TNBS group).

We next examined the small bowel but we did not find any overt inflammatory lesion in the duodenum or ileum (Figure 2*F* and *G*) despite increased TNF-α, IFN-γ and IL-12 proteins (Figure *2H*) and mRNA (Figure 2*I*) levels. As observed in CD patients, expression of *Nod2* was increased in the duodenum, ileum and the uninflamed part of the colon remote from rectal injury (Figure 2*I*). We hypothesized that this effect was consecutive to the recirculation of CD4^+^ pro-inflammatory T-cells in the gut mucosa. We therefore treated TNBS-challenged mice with anti-CD4^+^ monoclonal antibodies to reduce the number of CD4^+^ T-cells in the small bowel (Figure 2*J*). This treatment only partially improved the colitis but restored normal levels of *Nod2* and inflammatory cytokines in the duodenum and ileum (Figure 2*A-I*).

IFN-γ and TNF-α increase intestinal paracellular permeability via MLCK activation. We therefore investigated whether the paracellular permeability of the small bowel was affected in TNBS treated mice.^25^’^27^ Paracellular permeability as well as long *Mylk* mRNA expression were increased in both duodenum and ileum but returned to normal after CD4^+^ T-cell depletion (Figures 2*I* and *K*). Treatment of mice with an inhibitor of inflammatory CD4^+^ T-cells recirculation (FTY720) limited paracellular permeability increases in the duodenum and the ileum further indicating that a recirculation of T-cells from the rectal inflammatory lesions is likely responsible for the remote small bowel barrier loss (Figure 2*L*).

Of note, given the abundance of immune cells in inflamed areas, higher levels of epithelial NOD2 would be expected in the inflamed bowel of CD patients if the expression of epithelial *NOD2* was under the control of CD4^+^ T-cells. We therefore determined the populations of immune cells present in the lamina propria of CD patients by immunostaining. Consistent with data collected in mice, lamina propria CD4^+^ T-cell numbers were not increased in areas with the highest grades of inflammation. Most immune cells present in the lamina propria at these sites within the ileum (Figures 3*A* and *B*) and the colon (Figure 3*C* and *D*) were CD163+ macrophages. Consistently, CD4^+^ T-cells predominated in areas with low grade inflammation.

**Figure 3:**
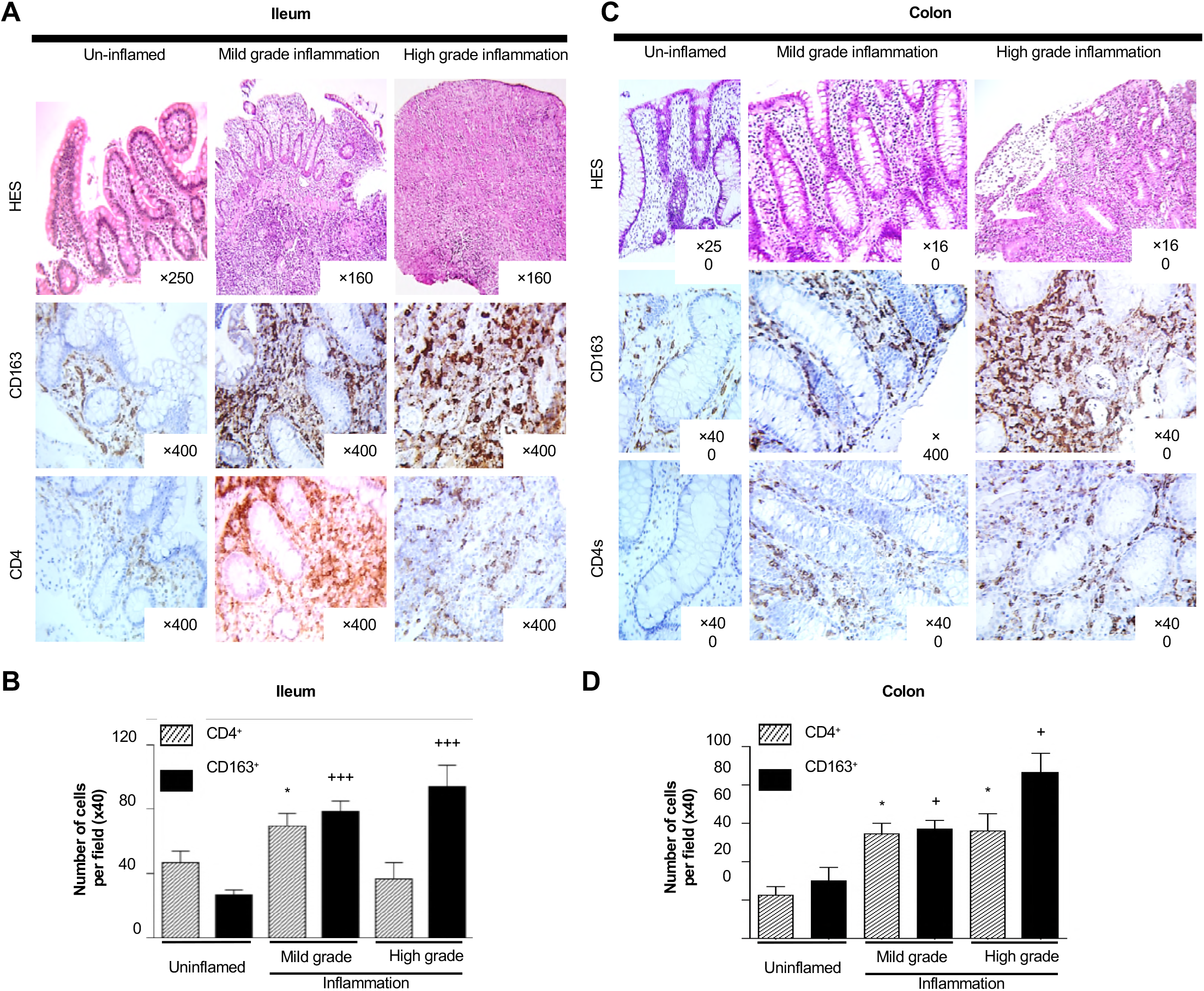
CD4 and CD163 immunostaining of ileal and colonic biopsies from CD patients. *(A-B)* Ileal and *(C-D)* colonic biopsies were collected from uninflamed or inflamed locations in CD patients. *(A* and *C)* Grading of the inflammation was confirmed by coloration by hematoxylin-eosin (HES) and CD4^+^ or CD163^+^ positive cells were assessed by immunostaining. *(B* and *D)* CD4^+^ or CD163^+^ positive cells were counted in the lamina propria. (At least n=6 fields per patients; mean ± SEM; *P<0.05 vs. uninflamed CD4^+^ T-cells; ^+^P<0.05 and ^+++^P<0.001 vs. uninflamed CD163^+^ T-cells). Areas with lymphoid follicles were excluded.

### MLCK activity is necessary to maintain the pro-inflammatory status of the small intestinal mucosa

Since pro-inflammatory cytokines such as IL-1β, TNF-α and IFN-γ can alter the paracellular permeability of the intestinal epithelium by increasing the expression and activity of long MLCK, we explored the role of MLCK in barrier function. ^17, 27–30^Treatment of mice with ML-7 (an inhibitor of MLCK) had only a limited impact on the severity of TNBS-induced colitis (Figure 4*A-E*). In contrast, MLCK inhibition restored normal TNF-α, IFN-γ, IL-1β, IL-12 expression (mRNA and protein), *Mylk* and *Nod2* mRNA transcription, and paracellular permeability in the duodenum and the ileum (Figures 4*F-H*). Similarly, knockout mice lacking long MLCK developed a slightly less severe colitis compared to WT mice (Figure 4 *I-L*) and did not develop increased duodenal or ileal paracellular permeability (Figure 4*M*). These data confirm that MLCK activity is responsible for the gut barrier defect remote from inflammatory lesions.

**Figure 4:**
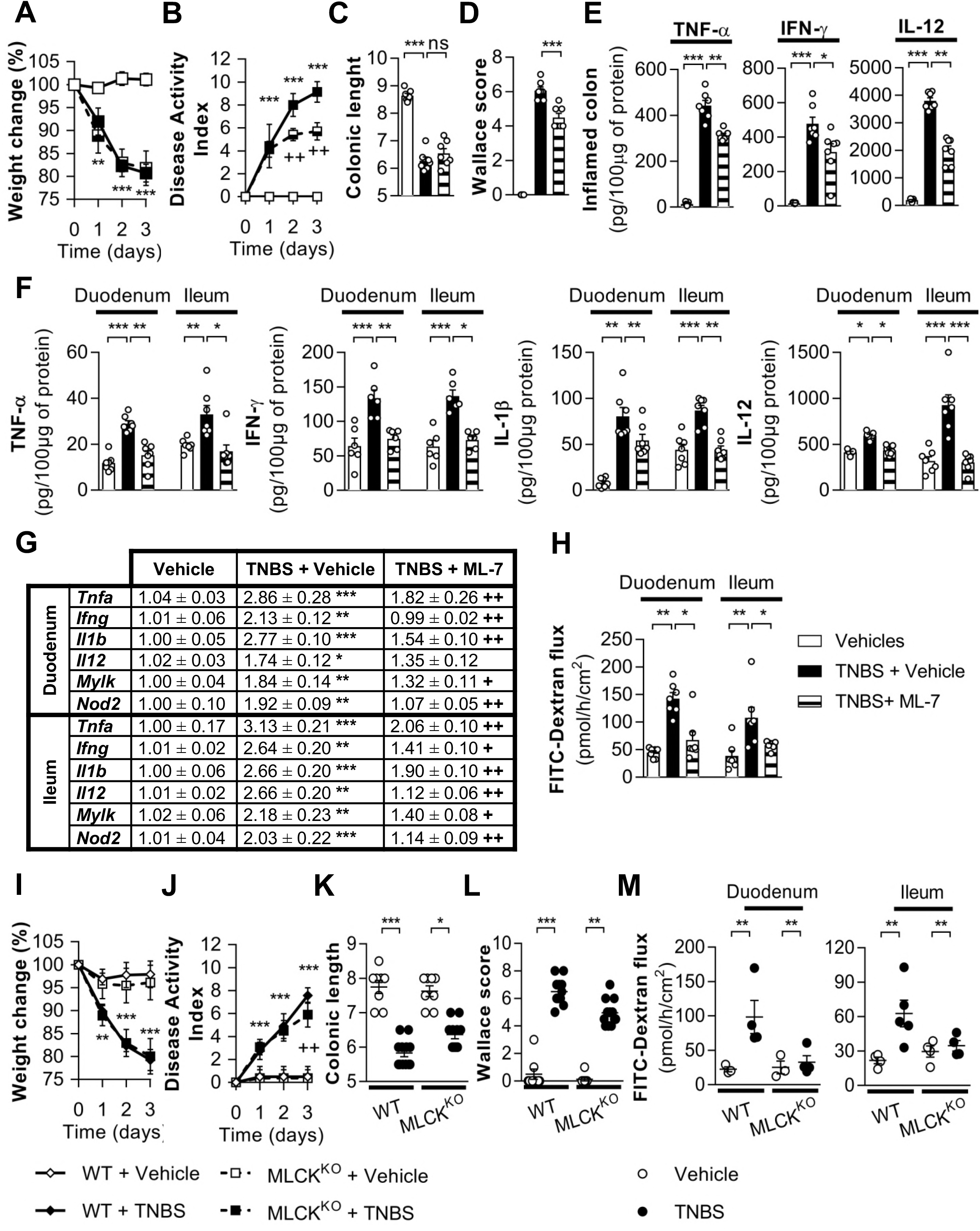
Inhibition of MLCK prevents the small bowel alteration triggered by TNBS induced colitis. *(A-M)* Colitis was induced by intra-rectal administration of TNBS in *(A-H)* wild-type (WT) mice or (I-M) long isoform MLCK knock-out (MLCK^KO^) mice while the control group was challenged with PBS-Ethanol. Mice were treated with ML-7, an MLCK inhibitor, or PBS (Vehicle) where indicated. 3 days after induction of colitis, the following parameters were monitored to evaluate the severity of the colitis: *(A,I)* Weight loss and *(B,J)* Disease activity index; *(C,K)* Colonic length; *(D,L)* Wallace score; (E) Levels of pro-inflammatory cytokine in inflamed colon. In parallel, the following parameters were measured in the duodenum and ileum: (F) protein levels and; (G) mRNA expression of pro-inflammatory cytokines, *Mylk* and *Nod2; (H,M)* Paracellular permeability. (One point = one mouse; mean ± s.e.m; 3 independent experiments; ^*^P<0.05, ^**^P<0.01 and ^***^P<0.001 vs. vehicle control group or indicated group; ^++^P<0.01 vs. TNBS group; ns=non-significant).

### NOD2 maintains the barrier integrity on remote small bowel

We have previously shown that stimulation of epithelial NOD2 with MDP allows the maintenance of the gut barrier.^17^ *Nod2*^WT^ mice were treated with MDP for 2 consecutive days before experimentation. Intraperitoneal injection of rhodamine-labelled MDP confirmed the ability of MDP to enter the enterocytes (Figures 5*A-C*). NOD2 stimulation reduced the disease activity index, colonic length, Wallace damage scores and pro-inflammatory cytokine expression without any effects on body weight loss after rectal TNBS infusion (Figures 5*D-H*). In contrast, in the small bowel, MDP treatment normalized mRNA (Figure 5*I*) and protein (Figure 5*J*) levels of pro-inflammatory cytokines as well as the paracellular permeability (Figure 5*K*). As expected, MDP did not affect the increase in *Nod2* expression (Figure 5*I*).

**Figure 5:**
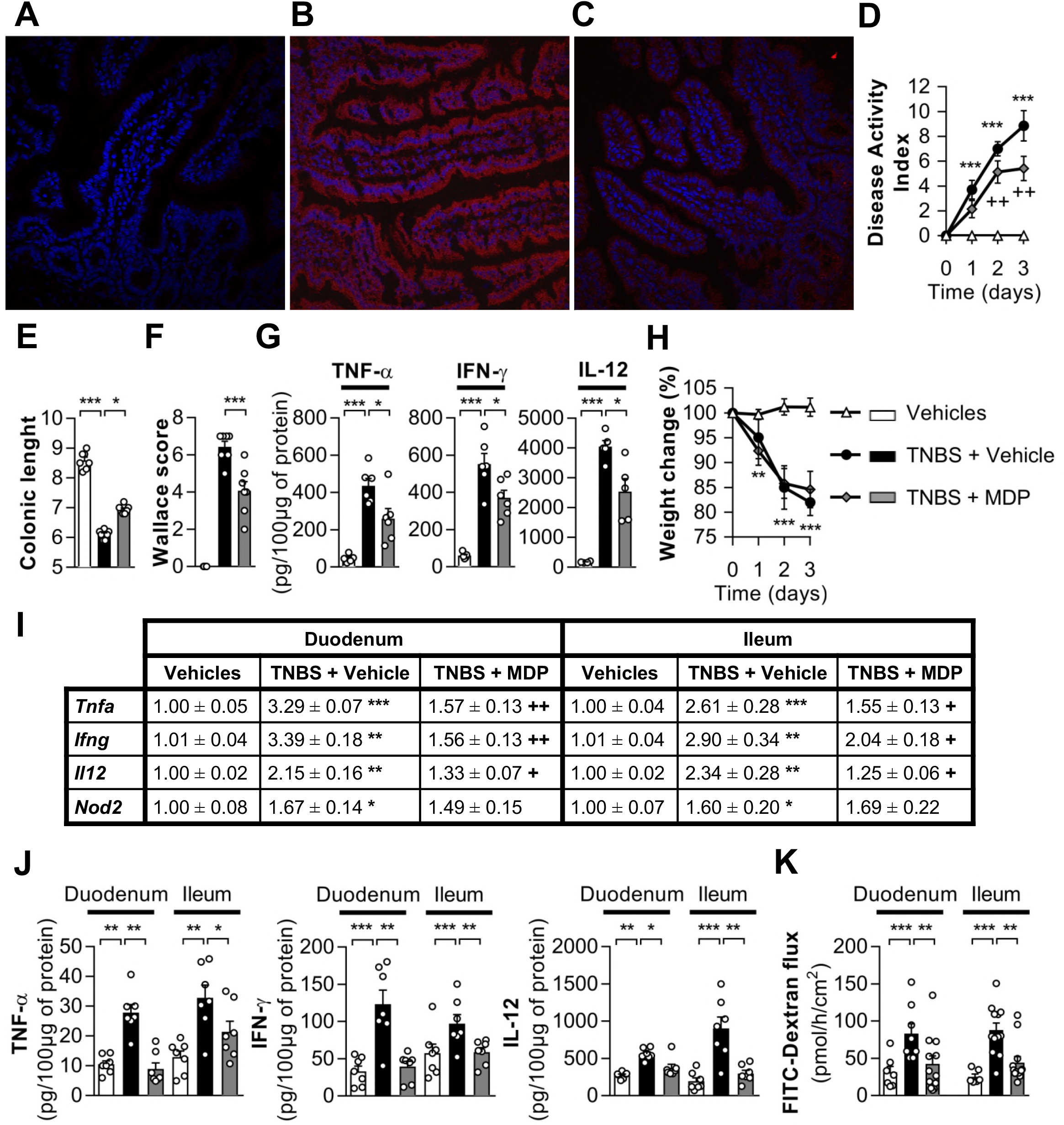
NOD2 activation reverses the remote effects of TNBS-induced colitis. *(A-C)* Localization of muramyl dipeptide (MDP) in the small intestine after intraperitoneal injection. Rhodamine-labeled MDP is detected in epithelial cells of the small intestine two hours after IP injection. Fluorescence (red) was detected in epithelial cells of the *(B)* duodenum and *(C)* ileum. Nuclei were stained with DAPI (blue). *(A)* The ileum of a mouse injected with distilled water was used as a negative control. Original magnification, X40. Scale bars: 100μM. *(D-K)* C57BL/6 wild-type mice were instilled intra-rectally with TNBS. Vehicle control group was challenged with PBS-Ethanol. Mice were treated with MDP where indicated. 3 days after induction, the colitis was monitored by the following parameters: *(D)* Disease activity index; *(E)* Colonic length; *(F)* Wallace score; *(G)* levels of pro-inflammatory cytokines in inflamed colon; *(H)* Weight loss. In parallel, the following measures were made in the duodenum and ileum: *(I)* mRNA expression of pro-inflammatory cytokines, *Mylk* and *Nod2; (J)* protein expression of pro-inflammatory cytokines; (K) Paracellular permeability. (Each point = one mouse; mean ± s.e.m; 3 independent experiments; ^*^P<0.05, ^**^P<0.01 and ^***^P<0.001 vs. vehicle group or indicated group; ^+^P<0.05, ^++^P<0.01 and ^++^P<0.001 vs. TNBS vehicle group).

Ablation of *Nod2* in mice (*Nod2*^KO^) results in increased paracellular and transcellular permeability across Peyer’s patches^14, 15^ and higher percentages of pro-inflammatory CD4^+^ T-cells^14^ but *Nod2*^Ko^ mice are only slightly more susceptible to TNBS-induced colitis (Figures 6*A-D*).^14^ However, while TNBS-treated *Nod2*^WT^ mice exhibit no lesion in the small bowel (Figure 2*F*), two thirds of *Nod2*^Ko^ mice showed overt duodenal inflammatory lesions as shown by a slight infiltration of scattered neutrophils in the *lamina propria* (Figure 6*E* and *F*). In the ileum, we observed a marked inflammation in 5/8 *Nod2*^ko^ mice, an infiltration of neutrophils and mononuclear cells in the villi and the crypts and a loss of muco-secretion. In addition, *Nod2^Ko^* mice exhibited an increased expression of pro-inflammatory cytokines in the duodenum and the ileum (Figure 6*G*). In contrast to *Nod2*^WT^ mice, treatment with MDP did not correct the expression of pro-inflammatory cytokines nor the increased permeability in the intestine of *Nod2*^Ko^ mice (Figures 7*A-F*). These findings indicate that the absence of *Nod2* leads to the development of remote lesions distant to rectal injury.

**Figure 6:**
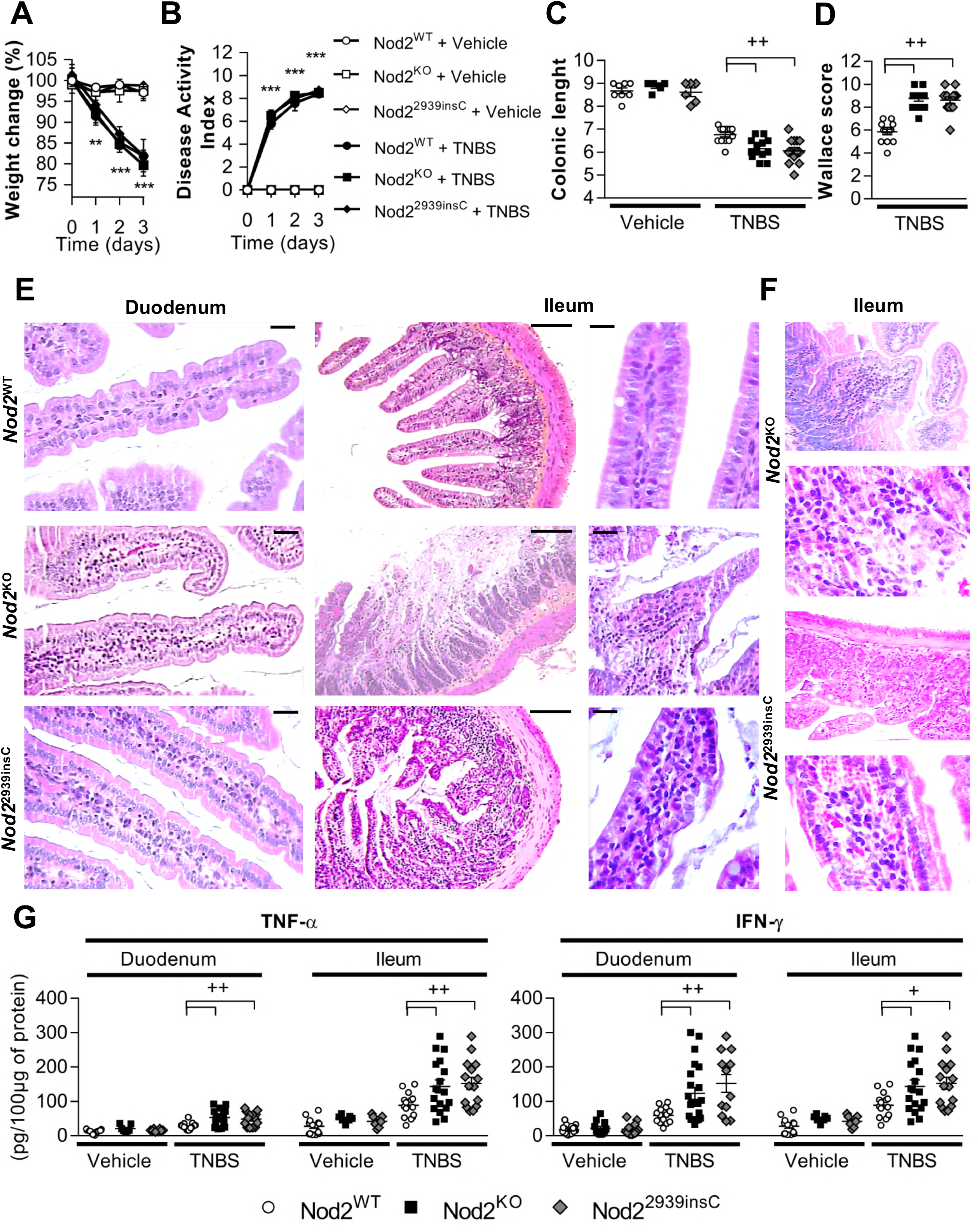
TNBS induced colitis leads to small bowel inflammation in *Nod2* deficient or mutated mice. *(A-F) Nod2*^WT^, *Nod2*^KO^ and *Nod2*^2939insC^ mice were challenged by intra-rectal instillation of TNBS. 3 days after induction, the colitis was monitored with the following parameters: *(A)* Weight loss; *(B)* Disease activity index; *(C)* Colonic length; *(D)* Wallace score. In parallel, the duodenum and ileum were studied by: *(E* and *F)* microscopic examination after hematoxylin-eosin staining; *(G)* protein levels of pro-inflammatory cytokines. (One point = one mouse; mean ± s.e.m; 3 independent experiments; ^**^P<0.01 and ^***^P<0.001 vs. Vehicle group; ^++^P<0.01 vs. indicated group; ns=non-significant).

**Figure 7:**
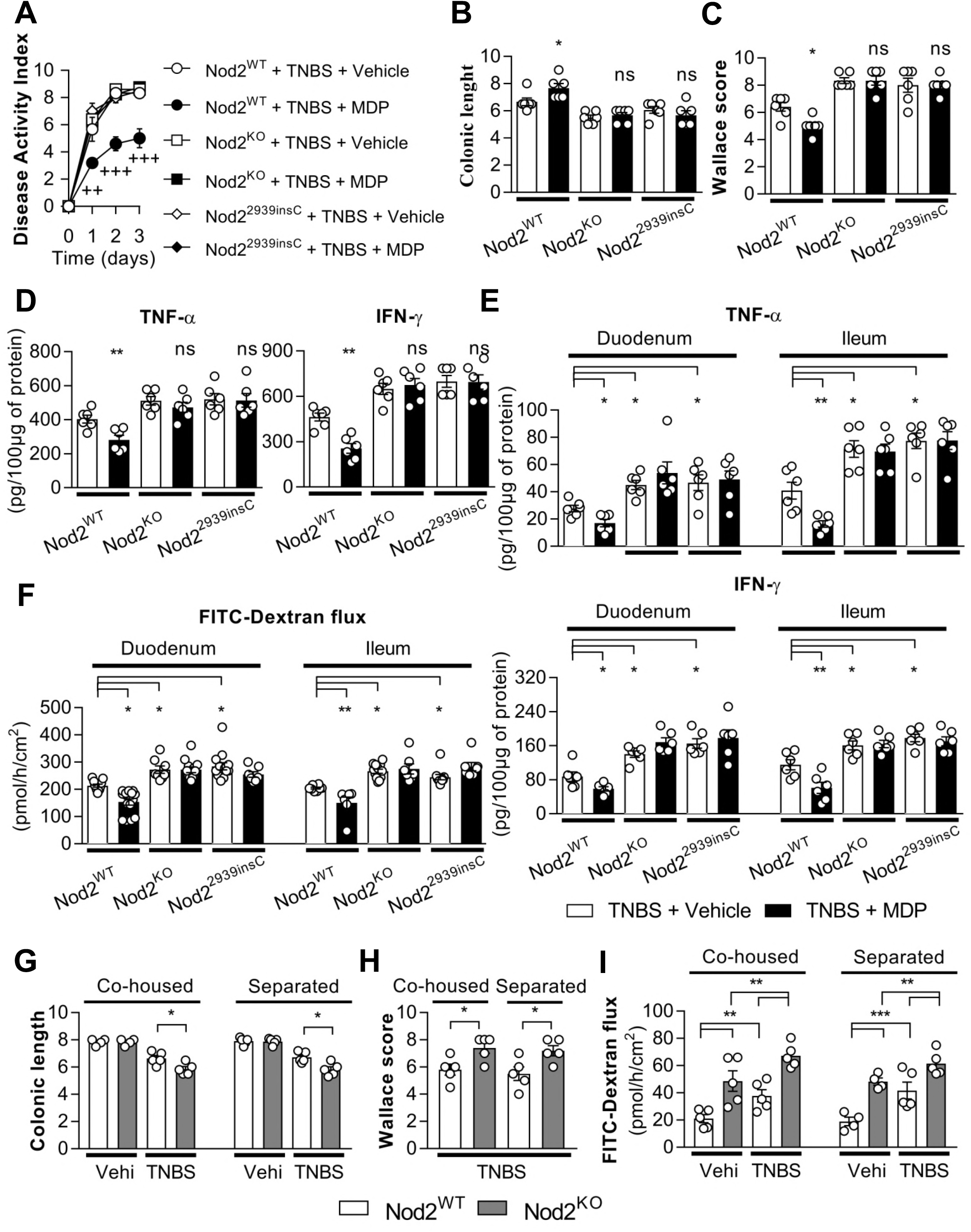
The small bowel is not protected by muramyl dipeptide in Nod2 deficient mice. The gut microbiota does not play a major role. *(A-E) Nod2*^WT^, *Nod2^KO^* and *Nod2*^2939insC^ mice were challenged by intra-rectal instillation of TNBS. Mice were treated with muramyl dipeptide (MDP) or PBS (vehicle) where indicated. *(A)* Disease activity index; *(B)* Colonic length; *(C)* Wallace score; Levels of pro-inflammatory cytokine in the (D) colon or (E) small bowel. (F) paracellular permeability in the small bowel.(G-I). *Nod2*^WT^ and *Nod2^*KO*^* mice were left separated or co-housed for at least 6 weeks to homogenize their microbiota. 3 days after TNBS-colitis induction, the following parameters were monitored: (G) Colonic length; (H) Colonic macroscopic score (Wallace score) and; (I) ileal Paracellular permeability (One point = one mouse; mean ± s.e.m; 3 independent experiments; ^*^P<0.05 and ^**^P<0.01 vs. indicated group; +P<0.05, ^++^P<0.05 and ^+++^P<0.001 vs. instilled TNBS *Nod2*^WT^ group).

In humans, CD is characterized by gastrointestinal skip lesions. Among the NOD2 genetic polymorphisms associated with CD, the 3020insC mutation encodes for a truncated (1007fs) protein. As described in *Nod2^ko^* mice, *Nod2*^2939insC^ mice-which are homozygotes for a mutation homologous to the Human 3020insC variant^12^-developed a slightly more severe colitis after TNBS administration (Figure *6A-D*). We observed inflammatory lesions in the duodenum and ileum of respectively 4/5 and 6/8 *Nod2*^2939insC^ mice after TNBS instillation (Figures 6*E* and *F*). Treatment of *Nod2*^2939insC^ mice with MDP did not reduce the expression of pro-inflammatory cytokines and the increased permeability in the small intestine (Figure 7*A-F*). We thus concluded that mice carrying a CD associated mutation of *NOD2* are not able to contain the intestinal inflammation where the primitive inflammatory lesions occurred.^12^

Since *Nod2*^Ko^ mice present a microbiota dysbiosis, we studied the contribution of gut microbiota in the inflammatory phenotype using littermate mice cohoused for 6 weeks.^31–34^ Sharing the dysbiotic microbiota associated with the deletion of *Nod2* in *Nod2* ^WT^mice did not change the severity of TNBS-induced colitis (Figures 7*G-H*) and the increased paracellular permeability of the ileal mucosa (Figure 7*I*). This finding suggests that the microbiota plays no major role on remote intestinal sites.

### Both hematopoietic and non-hematopoietic Nod2 regulate the small bowel function remote from colonic injury

NOD2 is detected in intestinal cells of hematopoietic and non-hematopoietic-origins.^35^ To compare the role of hematopoietic vs non-hematopoietic NOD2 in the small bowel during colitis, we compared *Nod2* chimeric mice after bone marrow stem cell (BMSC) transfer from *Nod2*^KO^ to *Nod2*^WT^ mice (KO→WT) and *Nod2*^WT^ to *Nod2^KO^* (WT→KO) to control mice transplanted with BMSC of the same genetic background (WT→WT and KO→KO) (Figures 8*A-C*).^32^ Chimeric mice were then challenged with TNBS three months after BMSC transplantation. Chimeric mice transplanted with *Nod2*^Ko^ BMSC were slightly more susceptible to TNBS-induced colitis than chimeric mice grafted with *Nod2*^WT^ BMSC (Figures 8*D-H*).^17^ Indeed, body weight loss, DAI and colonic length were similar between mice receiving WT (WT→WT and WT→KO chimeric mice altogether referred to WT→WT/KO mice) or *Nod2*^ko^ bone morrow (KO→KO and KO→WT chimeric mice referred to KO→KO/WT mice). However, the Wallace score, IFN-γ and TNF-α levels were higher in colonic inflamed mucosae of chimeric mice receiving *Nod2* ^ko^BMSC compared to mice receiving *Nod2*^WT^ BMSC (Figures 8*D-H*).

**Figure 8:**
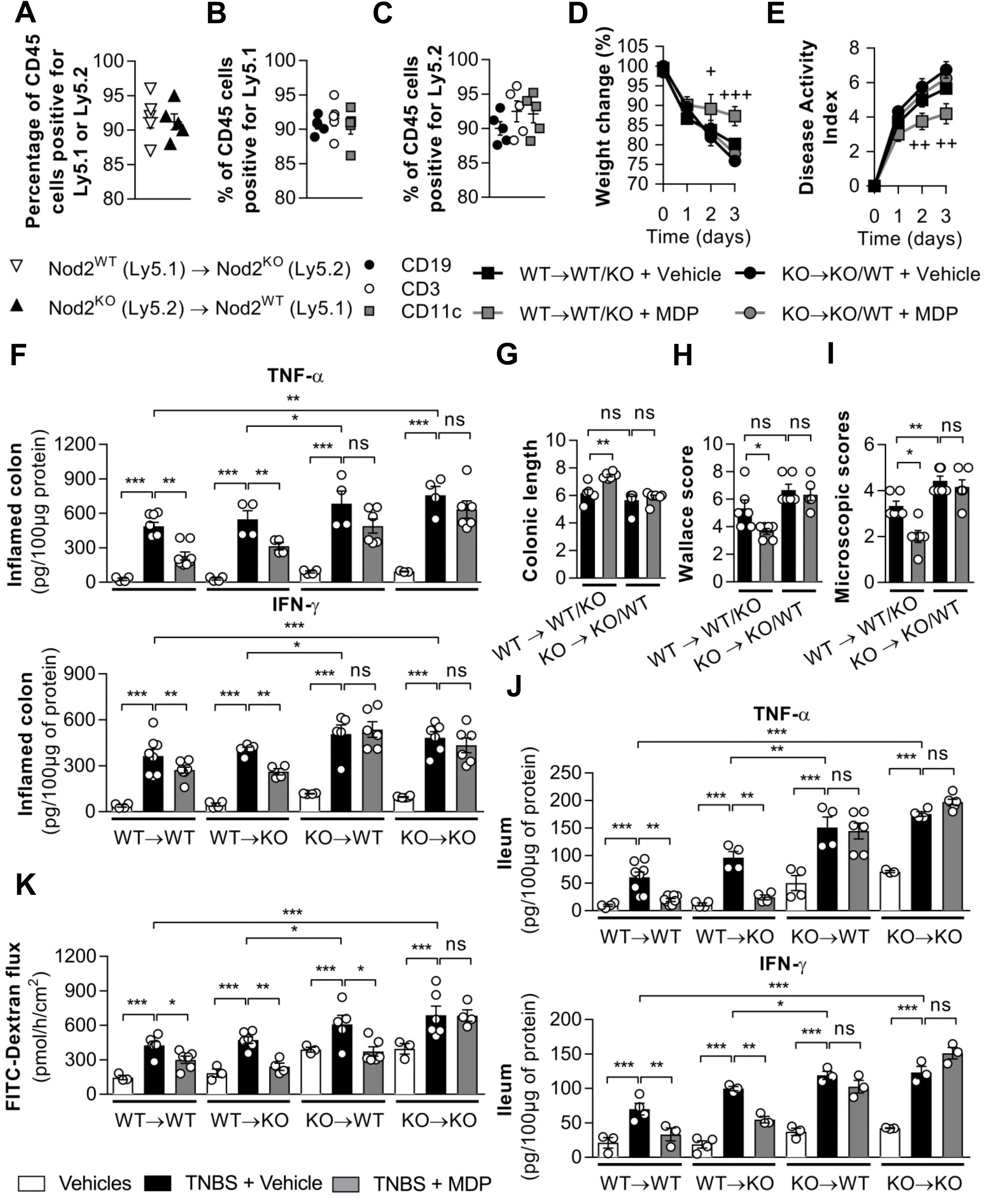
Both hematopoietic and non-hematopoietic *Nod2* regulate the small bowel function remote from gut injury. *(A-C)* Chimeric mice were generated by transplantation of BMSC mice from *Nod2*^WT^ to Nod2^KO^ (WT→KO) or from Nod2^KO^ to *Nod2*^WT^(KO→WT). Mice transplanted with BMSC of the same genetic background (WT→WT or KO→KO) served as controls. Where indicated WT→KO and WT→WT (respectively KO→WT and KO→KO) were pooled and annotated WT→WT/KO (respectively KO→WT/KO). Three months after bone marrow transplantation, chimerism for CD45 isoforms expression was monitored in Peyer’s patches of chimeric mice via flow cytometry. *(A)* Percentages of CD45.1 and CD45.2 positive cells. *(B* and *C)* Percentages of CD3+, CD19+ and CD11c+ cells in CD45Ly5.1 or CD45Ly5.2 respectively. *(D-K)* Three month after transplantation, colitis was induced by intra-rectal administration of TNBS. Mice were treated with MDP or PBS (vehicle) where indicated. 3 days after induction of colitis, the following parameters were monitored: *(D)* Weight loss; *(E)* Disease activity index; *(F)* Wallace score; *(G)* colonic length; *(H)* levels of pro-inflammatory cytokines in inflamed colon. In the ileum, the following parameters were recorded in parallel: *(I)* microscopic score; *(J)* Levels of pro-inflammatory cytokines and; *(K)* Paracellular permeability. (At least n=6 per group; mean ± s.e.m; data show a combination of two independent experiments; ^*^P<0.05, ^**^P<0.01 and ^***^P<0.001 vs. indicated group; ^+^P<0.05, ^++^P<0.01 and ^+++^P<0.001 vs. instilled TNBS control group; ns=non-significant).

Interestingly, ablation of *Nod2* in the hematopoietic lineages led to a more frequent and more severe inflammation in the ileum compared to chimeric mice expressing *Nod2* in the hematopoietic cells (Figure 8*I*). In parallel, expression levels of pro-inflammatory cytokines levels and paracellular permeability were higher in mice deficient for *Nod2* in BMSC (Figures 8*J* and *K*). Consistent with the anti-inflammatory role of NOD2 in the intestinal mucosa^36^, treatment with MDP improved the colonic inflammation but also the severity of the small bowel inflammation and the expression in inflammatory cytokines in the ileum only in chimeric mice expressing *Nod2* in their hematopoietic compartment (Figures 8*H-J*). However, chimeric mice expressing NOD2 in their radio-resistant compartment showed reduced paracellular permeability after MDP treatment regardless of the presence of NOD2 in hematopoietic cells (Figure 8*K*). This provides additional evidence that both hematopoietic and non-hematopoietic NOD2 exerts a protective function on the gut barrier.^17^

## Discussion

A “leaky gut” is a common feature of several conditions associated with *NOD2* mutations including CD. Here we show that NOD2 protects the small intestine not only in injured areas but also in areas remote from gut mucosal lesions. Indeed, NOD2 controls the paracellular permeability all along the digestive tract to contain the inflammation to local injuries and prevents its dissemination throughout the intestine.

We first observed that NOD2 expression was increased remote from primary inflammatory lesions in naïve pediatric CD patients. Interestingly, the increase in NOD2 expression was not restricted to immune cells in inflammatory areas as it was also detected in epithelial cells remote from CD lesions. We therefore hypothesized that epithelial NOD2 may have a specific role in healthy intestinal areas and explored the intestinal barrier remote from local injuries in mice.

The TNBS-induced colitis is a well-known model of self-limited inflammation. Although TNBS is administered in the rectum, it also alters the small intestine without any overt histological lesions in rats suggesting a remote effect of the colitis on the upper intestine.^24, 37^ In wild-type mice, we did not find any overt duodenitis or ileitis but we observed an increase in pro-inflammatory cytokines (TNF-α and IFN-γ) concentration, intestinal permeability and epithelial *Nod*2 expression in the small bowel. These effects were reversed by anti-CD4^+^ antibodies or inhibitor of recirculated CD4^+^ T-cells suggesting that they were consecutive to the recirculation of T-cells activated in the injured mucosa. Pharmacologic or genetic MLCK inhibition limited permeability increases, indicating that MLCK activation is a key component of the inflammatory response. Specifically, gut permeability augmented TNF-α and IFN-γ expression and altered lamina propria immune status.^21^ Conversely, pro-inflammatory cytokines increased paracellular permeability by stimulating MLCK expression and activity.^28, 29^

Local colonic injury leads to the disruption of the small bowel barrier. Since Nod2 is known to protect the gut barrier by inhibiting MLCK^15^ and because it was over-expressed in the small intestine, we supposed that it could restrain the leaky gut phenotype to the injured mucosa. MDP-induced activation of Nod2 fully corrected the impairment of the small bowel indicating that Nod2 plays a protective role along the small bowel. Recirculation of activated T-cells increases the gut permeability but also induces NOD2 expression which, in turn, strengthens the gut barrier. Interestingly, NOD2 seems to have little effect on the colitis itself. The severity of the inflammation may thus limit the effect of Nod2.

In contrast to WT mice, *Nod2*-deficient or mutated mice developed overt inflammatory lesions in the small bowel during TNBS-induced colitis, thus confirming the relevance of NOD2 in the protection of the gut barrier. Using BMSC transfer experiments (from *Nod2*^WT^ to *Nod2*^KO^ mice and vice-versa), we showed that both hematopoietic and non-hematopoietic NOD2 are necessary to protect the small bowel mucosa.

To the best of our knowledge, the role of NOD2 remote to colonic inflammation had never been demonstrated. In CD, Th-1 oriented CD4^+^ T cells appear to be key effectors of gut inflammation and NOD2 expression^38^ and most treatments (anti-inflammatory drugs, immune-suppressors and anti-TNF-α antibodies) target CD4^+^ T cells. For instance, TNF-α antagonists diminish the severity of the disease and restore the gut barrier function.^22, 39^ In our model, *Nod2* invalidation in the hematopoietic compartment is sufficient to promote a barrier defect which is consistent with reported cases of CD patients cured by allogenic or autologous hematopoietic stem cell transplantation.^40^ However, activation of epithelial NOD2 may also counteract the effect of IFN-γ and TNF-α suggesting that treatment of patients not carrying mutations in *NOD2* with NOD2 agonists could activate the negative feedback loop to prevent the propagation of the inflammation and the skip lesions defining CD.^41^

## Material and Methods

### Patients and biopsies

Intestinal biopsies were obtained from 17 untreated children during routine endoscopies performed to establish CD diagnosis. Controls were histologically normal digestive biopsies obtained from 5 children without inflammatory bowel disease. For each participant, one or two biopsies from the ileum and/or cecum were sampled. Biopsies were either immediately frozen and later stained with toluidine blue or fixed in 4%-phosphate-buffered formalin and stained with hematoxylin and eosin. All biopsies were graded histologically so that immunohistochemistry and laser microdissection could be correlated with disease severity. NOD2 immunostaining was performed as previously described with two different rabbit polyclonal antibodies (Cayman Chemical and a gift from G Thomas CEPH).^23^ Laser microdissection was performed on 7μm sections obtained from the frozen biopsies. After verification of the quality of tissues and the absence of ulcers, surface epithelial cells and lamina propria cells were laser-microdissected using a Leica^R^ AS LMD system (Leica microsystems) in less than one hour. A mean of 500 cells were microdissected from each of the specimens (range 100-1000 cells) and stored in Trizol^R^ reagent (Invitrogen, Groningen, The Netherlands). The study was approved by the ethic committee “de protection des personnes” (Saint Louis Hospital, Paris, France) and all the parents of participants provided a signed informed consent.

### Animal models

Housing and experiments were conducted according to institutional animal healthcare guidelines and were approved by the local ethical committee for animal experimentation (Comité Régional d’Ethique en matière dΈxpérimentation Animale no. 4, Paris, France). C57BL/6 wild-type (WT), *Nod2* null allele (*Nod2*^KO^) and *Nod2*^2939insC^ mice (homozygotes for a mutation homologous to the Human 3020insC variant) together with long *MLCK* deficient mice (MLCK^KO^) were generated or hosted in a pathogen free animal facility.^12, 14, 26^ The animal facility was monitored every six months in accordance with the full set of FELASA high standard recommendations. The putative impact of Nod2-related dysbiosis on the studied phenotypes was assessed using WT and *Nod2^KO^* mice cohoused for 6 weeks in the same cage where indicated.^33, 42^

For the construction of chimeric mice, five million bone marrow stem cells (BMSC) were isolated from WT Ly5.1 or *Nod2*^KO^Ly5.2 mice and injected intravenously either into WT Ly5.1 or *Nod2*^KO^ Ly.5.2 lethally-irradiated recipients. ^30,32^ Chimerism was verified at week 12 by flow cytometry using Ly5.1 and Ly5.2 congenic markers (Figure 8*A-C*).^35^

CD4^+^ T-cells were depleted by two intra-peritoneal (i.p.) injections of 100μg purified GK1.5 (anti-L3T4 (CD4^+^) monoclonal antibody (Pharmingen, Germany), 96 and 24 hours before experimentation and 24hours after TNBS administration.^15^ The effectiveness of CD4 + depletion in Peyer’s plates is shown in Figure 2*J*. To inhibit the recirculation of CD4^+^ T-cells, mice were treated i.p. with FTY720 (3mg/kg; Sigma, France)^43^ 0, 1, 2 and 3 days after TNBS infusion.

MLCK inhibition was achieved by ip injection of ML-7, 2 mg/kg body weight (Sigma, France) twice daily during 4 days before experiments and 24 hours after TNBS administration.^15^ To investigate the effect of Nod2 stimulation, adult mice were pre-treated i.p with muramyl dipeptide (MDP, 100μg/mice/day; Sigma, France) for 2 consecutive days before experimentation and 24 hours after TNBS administration.^35^

### Colitis induction

Colitis was induced in 12 weeks old mice by a single intra-rectal administration of 2,4,6-trinitrobenzene sulfonic acid (TNBS, Sigma, France), which was dissolved in ethanol (50:50 vol/vol) at a dose of 120 mg/kg body weight under anaesthesia. Groups used as controls (vehicle) received an equal volume of PBS and Ethanol (50:50 vol/vol) intra-rectally. A 100 μl aliquot of the freshly prepared solution was injected into the colon, 4 cm from the anus, using a 3.5 F polyethylene catheter as previously described.^14^ Body weight loss and disease activity index were monitored before and 72h after TNBS administration. Mice were sacrificed by cervical dislocation. Colonic length and macroscopic damage Wallace score were recorded.^44^

Duodenal and ileal samples were fixed in 4%-phosphate-buffered formalin and embedded in paraffin. Five-micrometer sections were cut and stained with hematoxylin and eosin. Grading of the inflammatory scores were performed in blind fashion according the follow criteria^45^: 0, no sign of inflammation; 1, very low level of leukocyte infiltration; 2, low level of leukocyte infiltration; 3, high level of leukocyte infiltration, high vascular density, and thickening of the colon wall; 4, transmural infiltration, loss of goblet cells, high vascular density, and thickening of the colon wall in less than half of circumference; 5 necrosis of more than half the circumference and transmural inflammation.

Myeloperoxydase (MPO) expression was detected by immunohistochemistry. All sections were deparaffinized in xylene, rehydrated, incubated in 3/ hydrogen peroxide for endogenous peroxidase removal, and heated for 10 minutes in sub-boiling 10 mM citrate buffer (pH 6.0) for antigen retrieval. Then, sections were processed using the ImmPRESS polymer detection systems & reagents (Vector Laboratories, Burlingame, Ca), using anti-MPO antibody (Abcam, Cambridge, UK).

### Muramyl dipeptide localization

Mice were injected intraperitoneally with 300μg of rhodamine-labeled muramyl dipeptide (MDP, InvivoGen, San Diego, CA). Two hours later, mice were anesthetized with isofurane (Centre Specialités Pharmaceutiques, Moussy-le-Neuf, France) and sacrified. Ileal and duodenal samples were collected and rinsed with ice-cold PBS (ThermoFisher, Waltham, MA). Tissue was frozen in liquid nitrogen using HistoLab OCT cryomount (Histolab, Gothengurg, Sweden), 10μm-thick cryosections were cut and then fixed in 4%. PFA. MDP-rhodamine localization was detected by fluorescence confocal microscopy (confocal sp8, Leica, Frankfurt am Main, Germany).

### Paracellular permeability measurement

To measure the intestinal permeability, biopsies from duodenal and ileal mucosa were mounted in a Ussing chamber exposing 0.196 cm^2^ of tissue surface to 1.5ml of circulating oxygenated Ringer solution at 37°C. Paracellular permeability was assessed by measuring the mucosal-to-serosal flux of 4 kDa FITC-dextran (Sigma, France).^30^

### ELISA

Biopsies of duodenum, ileum and colon from different mice models were collected and washed with cold PBS. These biopsies were then homogenized using an ultra-thurax in 1 ml of PBS1X and, the concentration of protein was determined using commercial kit (Biorad, Marnes la Coquette, France). IFN-γ, IL-1β, IL-12 and TNF-α protein levels in the intestine were determined by ELISA according to manufacturer’s instructions (BD Biosciences).^46^

### DNA extraction and real time quantitative PCR

After extraction by the NucleoSpin RNA II Kit (Macherey-Nagel, France), total RNAs were converted to cDNA using random hexonucleotides and then used for RT-PCR (Invitrogen). We conducted qPCR with QuantiTect SYBR Green PCR Kit (Applied, France) using sense and antisense primers specific for G3PDH, the long MLCK isoform (specifically expressed by epithelial cells), *Ifn-γ, Il-1β, Il-12, NOD2, Mylk* and *Tnf-α* (primers used available in table 1). The cycle threshold (Ct) was defined as the number of cycles at which the normalized fluorescent intensity passed the level of 10 times the standard deviations of the baseline emission calculated on the first 10 PCR cycles. Results are expressed as 2^-ΔΔCt^ as previously described.^33^ For RNA samples obtained by laser microdissection, NOD2 expression was measured in triplicate and normalized using the Abelson housekeeping gene. To derive a relative number of mRNA molecules, a titration curve was established with NOD2 plasmids (from 1 to 10^6^ copy/microliters).

**Table 1.**
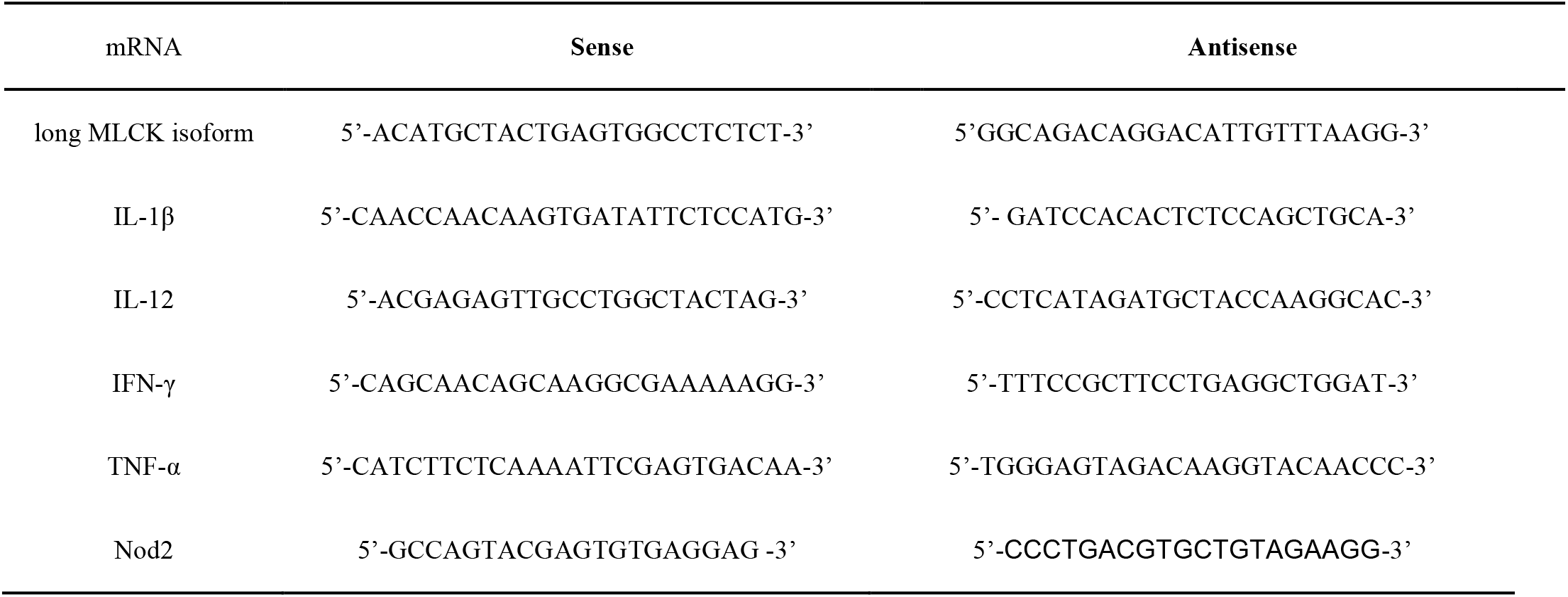
List of primers used for qPCR analyses in mice.

### Statistical analysis

For all the analysis, multigroup comparisons were performed using one-way ANOVA statistics with Bonferroni correction for multiple comparisons where an unpaired t-test assuming the Gaussian distribution was applied. The Gaussian distribution was tested by the Kolmogorov-Smirnov test. Statistical analyzes were performed using GraphPad Prism 7.00 (GraphPad Software). A two-sided P-value < 0.05 was considered statistically significant. All authors reviewed the data and approved the final manuscript.

## Acknowledgements

We thank Latifa Ferkdadji, Michel Peuchmaur, Xavier Fund, Anh Thu Gaston, Guillaume Even and Céline Berraud for their support and assistance. Financial support was provided by INSERM, Université Paris Diderot, Assistance Publique Hopitaux de Paris, Association François Aupetit and Investissements d’Avenir programme ANR-11-IDEX-0005-2, Sorbonne Paris Cite, Laboratoire d’excellence INFLAMEX and NIH grant NIDDK DK061931/DK068271.

## Author contributions

Study design and concept: ZA, DB, HZ, FB, JPH; Data acquisition: ZA, DB, CMV, NM, CM, MR, MD, HZ, CJ, FB; Analysis and interpretation: ZA, DB, CMV, NM, GD, JRT, CM, MR, NCB, CJ, FB, JPH; Writing of the manuscript: ZA, GD, JRT, NCB, FB, JPH; Obtained funding: JPH; Technical support: DB, CMV, GS, GD, NM, CM, MR, EOD, MD, CJ; Study supervision: DB, FB, JPH.

Authors have no conflict of interest to declare

## Abbreviations

CD: Crohn’s disease
IM: intestinal mucosa (without Peyer’s patches)
MDP: Muramyl dipeptide
MLCK: myosin light chain kinase
NOD2: nucleotide oligomerization domain 2
RICK: receptor-interacting serine/threonine kinase
TAK1: transforming growth factor β-activated kinase 1
TNBS: 2,4,6-trinitrobenzene sulfonic acid
TNF-R: TNF receptor
WT: wild-type

